# A Brain to Spine Interface for Transferring Artificial Sensory Information

**DOI:** 10.1101/807735

**Authors:** Amol P. Yadav, Daniel Li, Miguel A. L. Nicolelis

## Abstract

Lack of sensory feedback is a major obstacle in the rapid absorption of prosthetic devices by the brain. While electrical stimulation of cortical and subcortical structures provides unique means to deliver sensory information to higher brain structures, these approaches require highly invasive surgery and are dependent on accurate targeting of brain structures. Here, we propose a semi-invasive method, Dorsal Column Stimulation (DCS) as a tool for transferring sensory information to the brain. Using this new approach, we show that rats can learn to discriminate artificial sensations generated by DCS and that DCS-induced learning results in corticostriatal plasticity. We also demonstrate a proof of concept brain-to-spine interface (BTSI), whereby tactile and artificial sensory information are decoded from the brain of an “encoder” rat, transformed into DCS pulses, and delivered to the spinal cord of a second “decoder” rat while the latter performs an analog-to-digital conversion during a tactile discrimination task. These results suggest that DCS can be used as an effective sensory channel to transmit prosthetic information to the brain or between brains, and could be developed as a novel platform for delivering tactile and proprioceptive feedback in clinical applications of brain-machine interfaces.

## Introduction

Brain machine interfaces (BMI) have shown considerable promise as the basis of a new generation of assistive and restorative technologies for people suffering from severe neurological impairment, due to chronic spinal cord injuries and other devastating motor disorders^1, 2^, for a comprehensive review see^3^. Traditionally, BMIs transform raw electrical neuronal activity into motor commands that can control the real-time movements of artificial actuators like cursors^4^, robotic devices^2, 5^, and prosthetic limbs^6, 7^. Although BMI studies focused on ways to compensate for loss of motor function, proper somatosensory feedback remains a key component for the optimal operation of a BMI, particularly during clinical implementations, since such a feedback signal is essential for patients to achieve optimal BMI control and to incorporate artificial actuators, controlled by a BMI, as part of the patient’s body schema ^8, 9, 10^.

In the past, electrical intracortical microstimulation (ICMS) of the somatosensory cortex and stimulation of thalamic nuclei have been proposed as potential target sites for input of prosthetic sensory information^11, 12, 13, 14, 15, 16, 17, 18^; however, given the invasiveness of surgical procedures needed to implement these approaches in humans, alternative targets along the somatosensory pathway need to be explored for transmission of artificial sensory information to the brain.

Spinal cord stimulation (SCS) is an FDA-approved therapy for chronic pain. During the last decade, it has also emerged as a potential new therapy for treating freezing of gait, a notoriously difficult to treat manifestation of Parkinson’s disease^19, 20, 21^. SCS’s potential role in rehabilitation research has been explored only recently^22, 23^. While several studies have demonstrated the ability of SCS to repair injured spinal locomotor circuits^24, 25, 26^, raising the hypothesis that it could provide some type of functional and neurological recovery to chronic spinal cord injury patients, the potential role of SCS in directly modulating cortical circuits or enabling cortical plasticity remains poorly understood.

During the past decade, our laboratory has demonstrated in a series of studies in rodents and monkeys that continuous electrical stimulation of the dorsal columns of the spinal cord, a procedure called dorsal column stimulation (DCS), leads to modulation of supraspinal circuits in animal models of Parkinson’s disease and chronic epilepsy that is refractory to treatment^27, 28, 29, 30^. Based on these studies, we put forward the hypothesis that DCS could be used as a preferential pathway through which somatosensory feedback, generated by BMI-controlled artificial actuators, could be delivered to higher brain structures, such as the neocortex. In addition, we proposed that DCS could also be used to mediate the transfer of sensory information between multiple brains, an approach our lab pioneered and which we named a brain-to-brain interface (BTBI)^31, 32, 33, 34^.

In the present study, therefore, we substitute DCS for ICMS in order to test both hypotheses and in the process create a new approach which we named brain-to-spine interface (BTSI). To achieve this goal, though, first, we had to investigate whether rats can discriminate artificial sensations induced by DCS. To test that, we employed the traditional framework of reinforcement learning using a multiple alternative forced choice task. Secondly, if rats could learn to consciously differentiate artificial somatosensory stimuli generated by time-varying patterns of DCS, it was important to inquire whether this learned behavior resulted in corticostriatal plasticity. Finally, we studied whether DCS could be utilized for transmission of sensory information between two rat brains by pairing an encoder rat with a decoder rat and analyzing the performance of a BTSI.

## Results

### Learning to discriminate artificial somatosensory stimuli generated by DCS

We first investigated whether spinal cord electrical stimulation could be used to generate artificial sensations in adult rats (Fig. 1a). To this end, we trained animals on a multiple alternative forced choice task using a modified aperture-width discrimination chamber, created and perfected in our lab over the past 16 years. A full description of this task and its apparatus can be found elsewhere^35, 36^. In our modified task, rats had to discriminate artificial sensations induced by microstimulation pulses delivered to their spinal cord’s dorsal columns (Fig 1b and see Methods). In the two alternative version of this task [Pattern 1: 100 pulses at 333 (or sometimes 100 Hz: see Methods) vs Pattern 2: 1 pulse], rats’ discrimination performance started around chance (50 percent), and gradually increased from 50.83 ± 2.21 percent at session 1 to 91.75 ± 1.36 percent at session 11 (Fig 2a, n=12). This discrimination performance turned out to be comparable to how animals performed tactile discrimination of mechanical stimuli delivered to their facial whiskers^36, 37^. Rats learned to discriminate Pattern 1 stimuli at a slightly faster rate than they learned to discriminate Pattern 2 stimuli (3.57 ± 0.21 vs 2.46 ± 0.47, slope of blue vs red line in Fig 2a, p<0.05, linear regression, mean ± s.e.m.).

**Figure 1:**
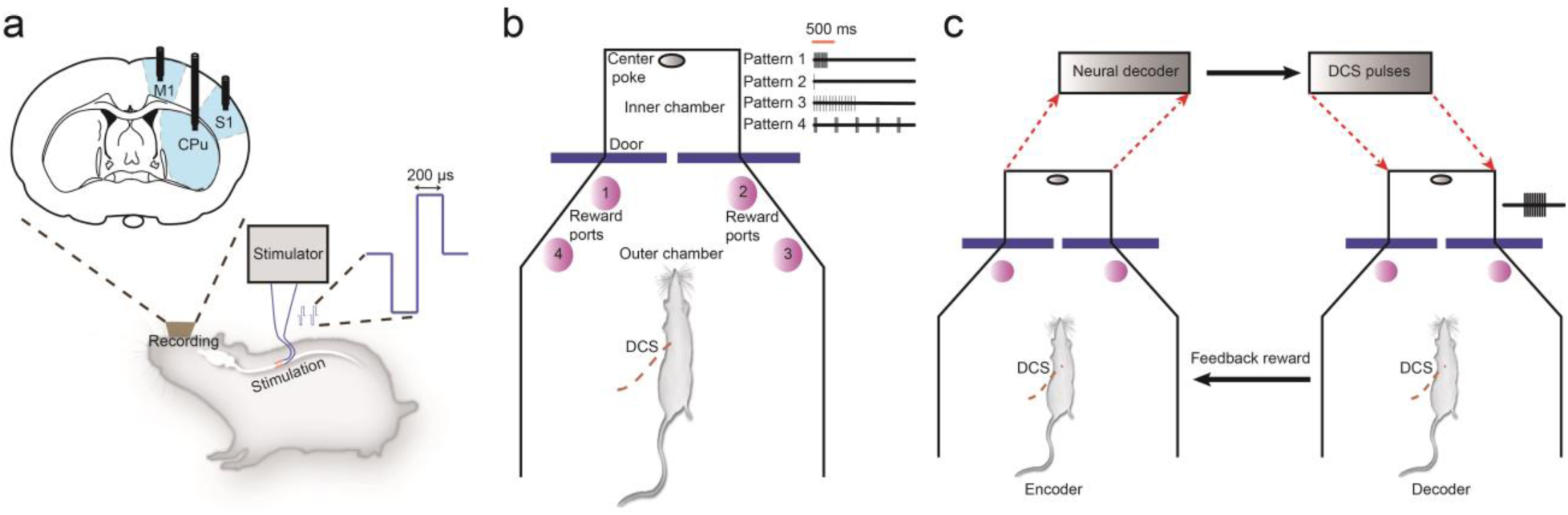
Experimental setup for artificial sensory discrimination using DCS and brain-to-spine interface. a) Rats were implanted with recording electrodes in motor cortex (M1), somatosensory cortex (S1) and striatum (STR) and dorsal column stimulating electrodes in the thoracic epidural space. Stimulation consisted of charge-balanced, biphasic pulses of 200 – 1000 µsec pulse duration delivered through an electrical microstimulator controlled by custom Matlab scripts. **b)** Behavioral setup for artificial sensory discrimination using DCS consisted of a modified aperture width tactile discrimination box where rats waited in the outer chamber until the sliding door opened. Then they had to poke into a hole in the center of inner chamber. Inside the inner chamber, depending on the type of trial, stimulation pulses were delivered. As the rats came out from the inner chamber they had to poke in the correct reward port associated with the stimulation pattern to get a water reward. **c)** Setup for the brain-to-spine interface consisted of two modified aperture width tactile discrimination boxes. While the encoder received DCS pulses inside the inner chamber on its way to the center poke, its post-stimulus neuronal activity was sent to a neural decoder (an adaptive logistic regression classifier) which predicted the number of DCS pulses (between 101 and 1) to be delivered to the decoder’s spinal cord. For the encoder, ‘virtual narrow’ width was rewarded with a response in the ‘left’ reward port while ‘virtual wide’ width was rewarded in the ‘right’ reward port. If the decoder chose the corresponding left or right reward port, based on the translated DCS pulses, the encoder received an additional feedback reward.

**Figure 2:**
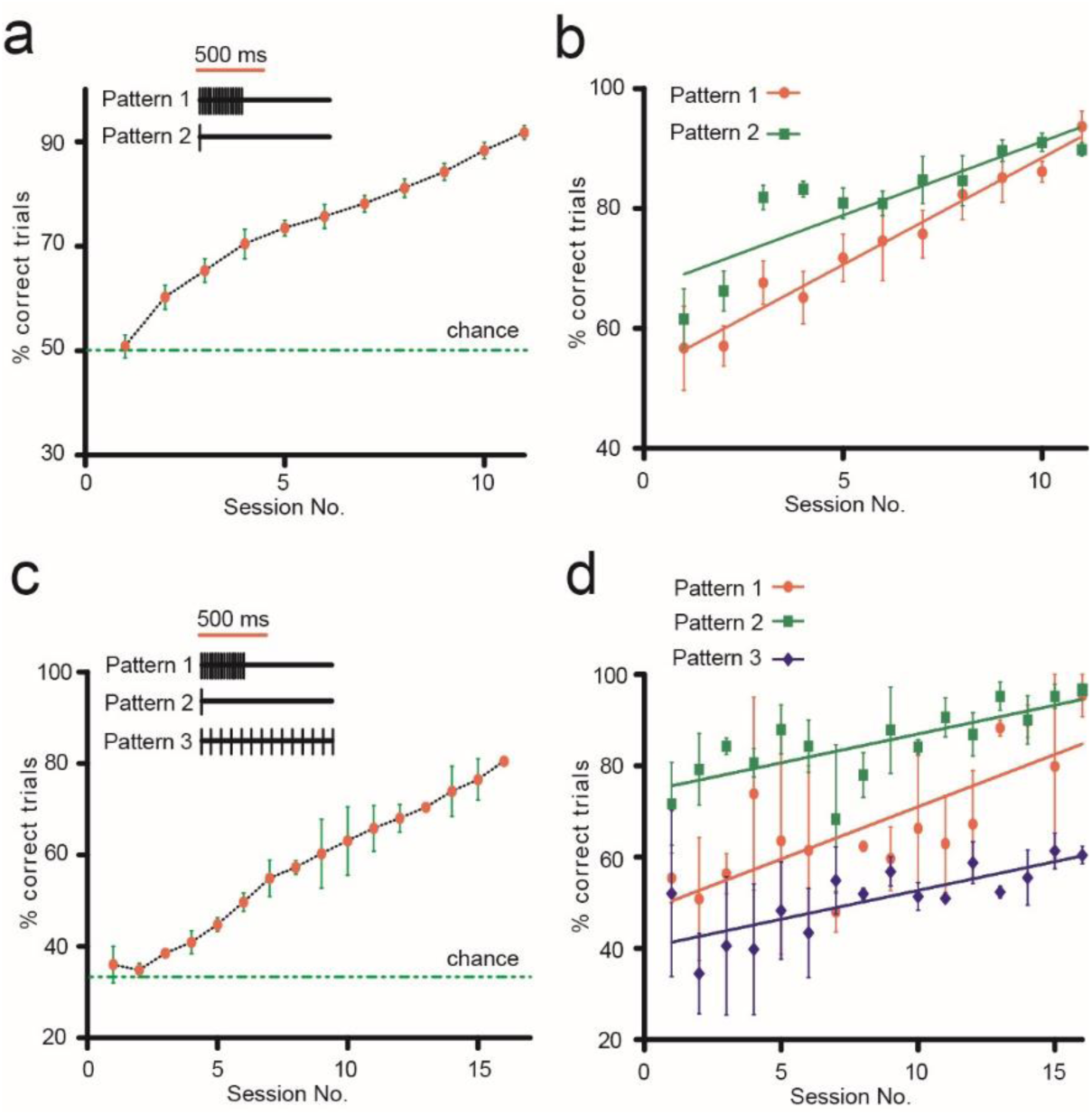
Rats learn to discriminate artificial sensations generated using DCS. a) Learning curve showing that rats learned to discriminate between two DCS patterns (Pattern 1:100 pulses and Pattern 2: 1 pulse). Graph shows percentage of correct trials as a function of session number (Red dots/green bars indicate mean ± s.e.m for 11 sessions in 12 rats, green line represents chance level of 50 %). **b)** Discrimination performance for each of the two patterns shown individually across sessions (mean ± s.e.m). **c)** Learning curve showing rats that learned to discriminate three DCS patterns (Pattern 1: 100 pulses at 333 Hz, Pattern 2: 1 pulse and Pattern 3: 100 pulses at 100 Hz). Graph shows percentage correct trials (Red dots/green bars indicate mean ± s.e.m for 17 sessions in two rats, green line represents chance level of 33 %). **d)** Discrimination performance for each of the three patterns shown individually across sessions (mean ± s.e.m).

Next, we explored whether rats could discriminate additional DCS stimulation patterns, if those were associated with a new reward port, i.e. (port 3 in Fig 1b). Now, rats that had previously learned to discriminate 100 pulses at 333 Hz vs 1 pulse learned to associate a novel stimulation pattern (Pattern 3: 100 pulses at 100 Hz), using a unique training paradigm (Supplementary Fig 1a and Methods), where task complexity was gradually increased. Two rats successfully learned to discriminate between these three patterns of stimulation (Fig 2c) reaching 80% accuracy in ∼15 sessions with max performance of 81%. The learning rate for either of the stimuli was not significantly different from the other (Pattern 1: 2.29 ± 0.61, Pattern 2: 1.26 ± 0.33, and Pattern 3: 1.26 ± 0.39, slope of red, green and blue lines respectively in Fig 2d, p>0.05, linear regression).

One of the tested rats went even further and learned to associate yet another stimulation pattern (Pattern 4, involving 5 bursts of 20 pulses each, with inter-burst frequency of 2 Hz and inter pulse frequency of 333 Hz, was associated with reward port 4, Supplementary Fig 1b). Subsequently, this rat learned to successfully discriminate all four patterns of stimulation, reaching 80.33 % performance in 22 sessions (Supplementary Fig 1c). The learning rates for individual stimuli during 22 sessions were significantly different. (Pattern 1: 0.13 ± 0.14, Pattern 2: 0.7 ± 0.27, Pattern 3: 1.68 ± 0.48, Pattern 4: 2.14 ± 0.25, slope of red, green, blue and gray lines in Supplementary Fig 1d respectively, p<0.0001, linear regression). Supplementary Figures 1e and 1f showcase how the percentages of response choices associated with each of the stimulation patterns gradually increased throughout the learning period, as rats learned to discriminate three and four stimulation patterns.

In addition to studying the behavioral latency, for changes between early (first) and late (last) training sessions, we investigated whether DCS induced learning was driven by cortical and striatal plasticity. This involved the analysis of local field potential (LFP) signals and neuronal spiking activity from motor cortex (M1), somatosensory cortex (S1), and striatum (STR), in rats that successfully learned to discriminate Patterns 1 and 2 (Fig 3a). Initially, we observed a significant difference in behavioral discrimination performance between early (52.67 ± 2.23%) and late (88.08 ± 2.37%) training sessions (p<0.01, Wilcoxon signed rank test, two-tailed, Fig 3b, n=12). Behavioral latency significantly increased from 0.26 ± 0.04 to 0.33 ± 0.02 seconds (p<0.05, Wilcoxon signed rank test, two-tailed, Fig 3c) between early and late sessions for seven rats that were previously trained on the whisker discrimination task (latency data from two rats was not recorded). Three naive rats decreased their behavioral latency, however not significantly, from 0.42 ± 0.08 to 0.28 ± 0.04 seconds (n.s. p>0.05, Wilcoxon signed rank test, two-tailed, Fig 3c) between early and late sessions. Overall, there was no difference between LFP spectral power, between early and late sessions across brain areas, when we considered LFP records for the entire session (Fig 3d). However, when we examined the spectral power of LFPs during the pre-stimulus period (300 ms before stimulation onset), we noticed a clear increase in striatal spectral power in the low frequency bands, between early and late sessions, as the rats (n=5 with striatal implants) learned the discrimination task (Fig 3e, 3f and 3g). Specifically, the mean power in the delta range (1.5 – 4.5 Hz; early: −15.77 ± 1.28 dB vs late: −14.14 ± 1.07 dB) and the mean power in theta range (5 – 9.5 Hz; early: −16.69 ± 1.23 dB vs late: −15.05 ± 1.03 dB) was significantly higher in late learning than early learning (Fig 3f and 3g, p<0.05, paired t-test, two-tailed). S1 and M1 LFP spectral power showed no statistically significant difference between early and late sessions (data not shown). Interestingly, the magnitude of the change in behavioral latency was strongly correlated with the magnitude of change in the delta band (r^2^=0.86, p<0.05, Pearson correlation, two-tailed, n=5) and theta band (r^2^=0.82, p<0.05, Pearson correlation, two-tailed, n=5) striatal spectral power (Fig 3h, 3i) suggesting that pre-stimulus striatal spectral power modulation was a strong predictor of the change in behavioral latency. This finding suggests that learning to discriminate DCS patterns altered the behavior of our rats, thereby resulting in striatal plasticity, although no clear indication of cortical plasticity was observed in the LFPs.

**Figure 3:**
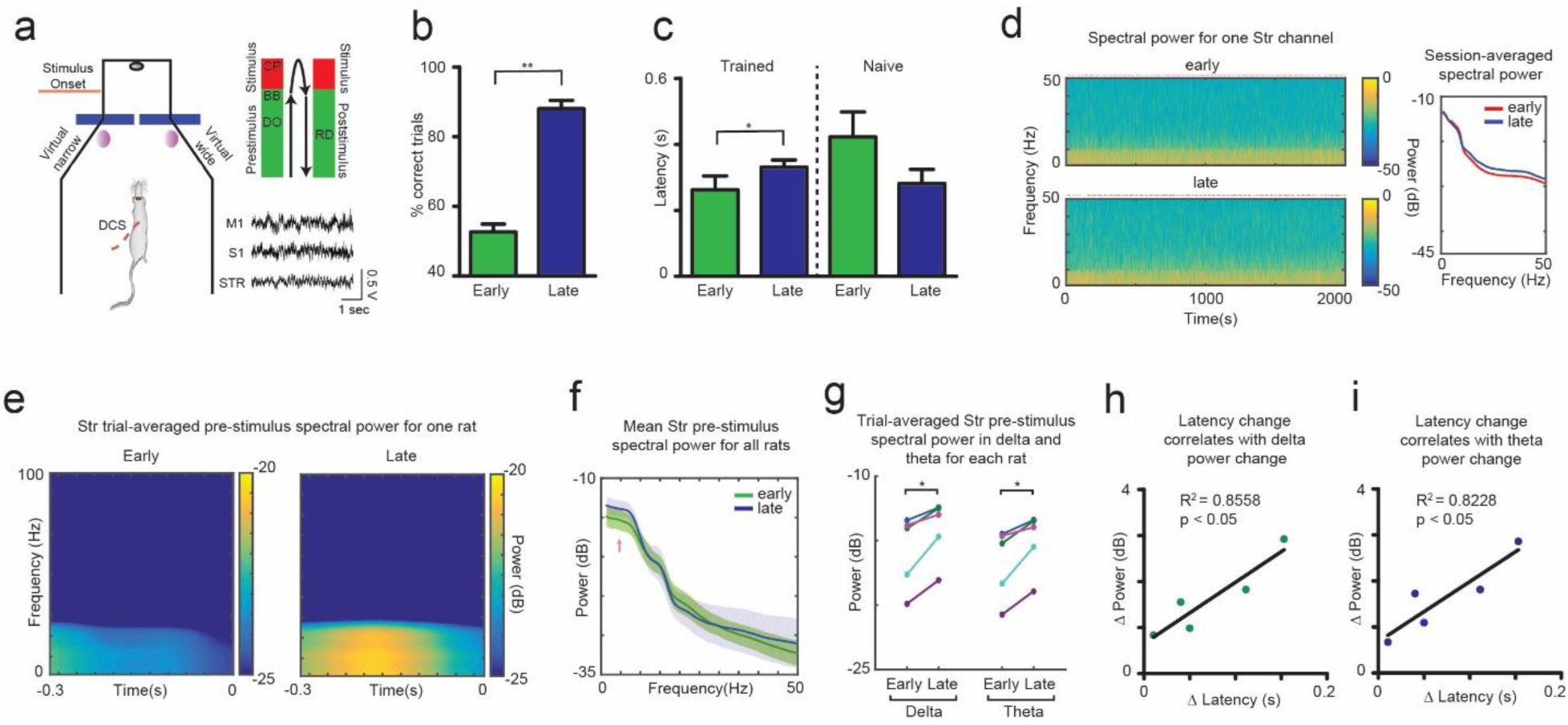
Learning to discriminate article sensations using DCS resulted in changes in striatal local field potentials (LFPs). a) Behavioral box used for training rats to discriminate two DCS patterns (Virtual narrow: 100 pulses vs. Virtual wide: 1 pulse). Each trial was divided into pre-stimulus, stimulus and post-stimulus periods. A typical trial comprised of the following sequence - DO: Door Open, BB: Beam Break, CP: Center Poke, RD: Reward delivery. Stimulation began at BB, when the rat crossed a photobeam inside the inner chamber. Spike and LFP activity from M1, S1 and STR was recorded during the training period and comparisons were made between early (1^st^ session) and late (last session) periods of training. **b)** Rats exhibited significant improvement in discrimination performance between early and late sessions. **c)** Behavioral latency (measured as time between BB and CP) significantly increased for rats (n=7) that were previously trained on the whisker discrimination task. Naive rats (n=3) decreased behavioral latency, however not significantly. **d)** Representative time-frequency spectrogram of same striatal LFP channel recorded during entire early and late session of one rat (red marker above graphs indicates stimulation onset time for each trial). Session-averaged spectral power for the same channel showing no difference between early and late sessions is shown at right. **e)** Trial-averaged pre-stimulus striatal LFP spectral power for one rat showing clear increase in the delta (1.5 - 4.5 Hz) and theta (5 - 9.5 Hz) frequency range from early to late learning. Time (0) corresponds to stimulation onset. **f,g)** Mean pre-stimulus striatal spectral power in the delta and theta range (pink arrow in f) significantly increased from early to late sessions in all rats (n=5)**. h,i)** Latency change between early and late sessions was significantly correlated with changes in pre-stimulus striatal delta and theta spectral power.

Although spiking activity from M1, S1, and STR areas was contaminated by the presence of stimulation artifacts during stimulation periods, such spurious signals were clearly distinguishable from actual spikes during offline sorting (Fig 4a, 4b). We corrected the spiking firing rate with artifact blanking (4 milliseconds, Fig 4c, 4d) and observed that there were clear differences in the firing rates of the neuronal ensemble between early and late sessions (Fig 4e, 4f). To quantify changes in neuronal ensemble firing patterns, we developed a custom adaptive logistic regression neural decoding classifier where binned firing rates were used as input features to predict the stimulus delivered on each trial. This adaptive algorithm updated its feature weights after each trial by adding the neuronal firing rates observed during previous trials to the training set, and using the updated dataset to re-train the classifier (a baseline training set was created using the first 20 trials of the session, see Methods for additional details). Thus, the feature weights were recalculated on each trial, based on firing patterns of all previous trials, and the current trial neuronal firing patterns were used for predicting the artificial sensory stimulus delivered to the rat’s dorsal columns in each trial. Prediction performance of the classifier, measured as a fraction of correct stimulus prediction and mean squared error (MSE), was significantly better in the adaptive version in comparison to the non-adaptive version – where the classifier was trained initially using the baseline training set and never updated (fraction correct: 0.91 ± 0.02 vs 0.82 ± 0.02, p<0.05, Wilcoxon signed rank test, two-tailed, n=6, Fig 4g; MSE: 0.07 ± 0.02 vs 0.14 ± 0.02, p<0.05, Wilcoxon signed rank test, two-tailed, n=6, Fig 4h). Performance of our custom classifier in predicting the artificial sensory stimuli was significantly better in late sessions, as compared to early sessions, in terms of both fraction correct predictions (early: 0.77 ± 0.05 vs late: 0.91 ± 0.02, p<0.05, paired t-test, two-tailed, n=6, Fig 4i) and MSE (early: 0.19 ± 0.04 vs late: 0.07 ± 0.02, p<0.05, paired t-test, two-tailed, n=6, Fig 4j). This suggests that as the rats learned to discriminate DCS patterns, their overall corticostriatal neuronal ensemble activity evolved to reflect an improved encoding of individual sensory stimuli. Similar changes in encoding of sensorimotor information by M1 ensemble activity were observed in an earlier study where rats learned a reaction-time motor task^38^.

**Figure 4:**
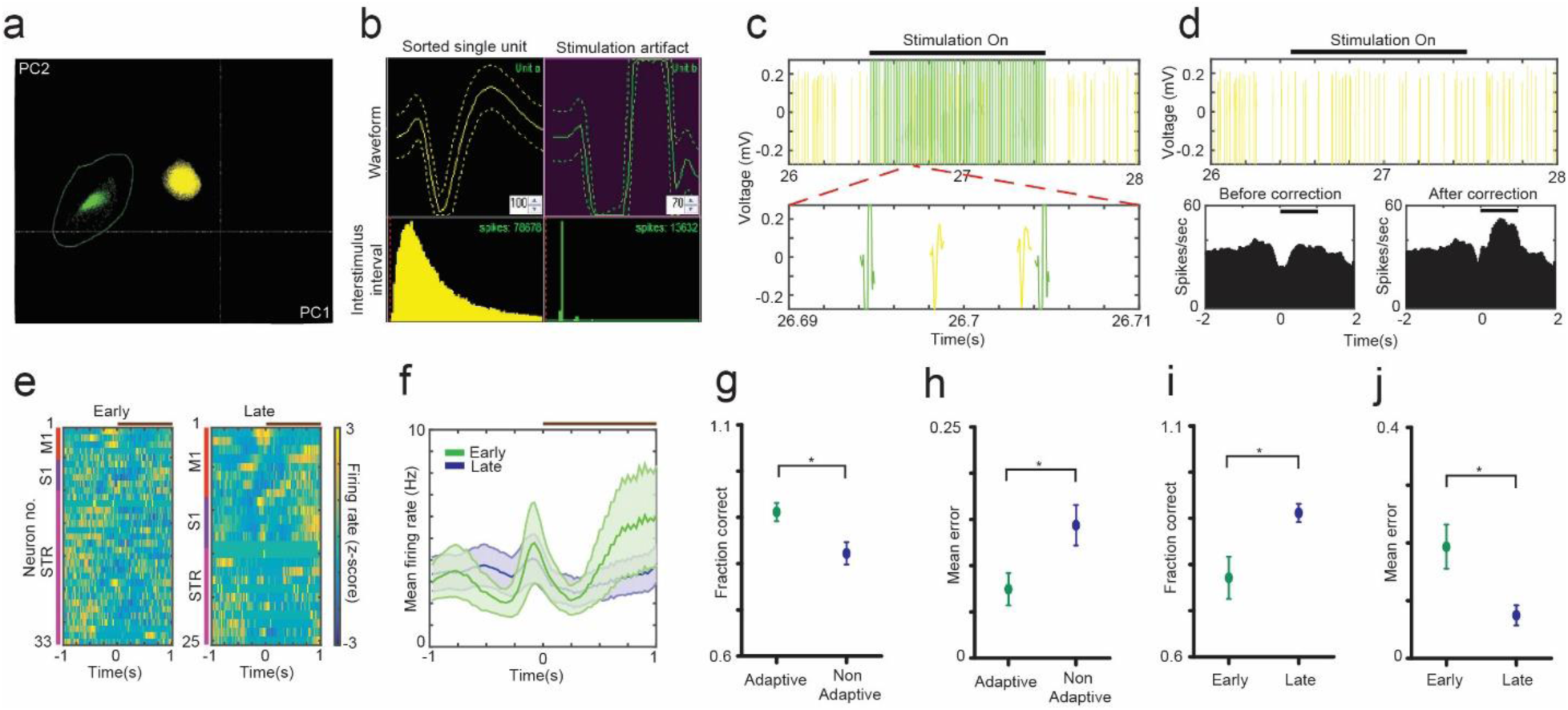
Learning to discriminate artificial sensations using DCS resulted in changes in M1, S1 and STR ensemble activity. a) Representative separation of single unit cluster from stimulation artifacts projected on the first two principle components during offline sorting. **b)** Single unit waveforms were clearly distinguishable from the stimulation artifacts. **c)** Stimulation waveforms resulted in amplifier dead time due to saturation during the ‘stimulation on’ period. A conservative blanking period of 4 msec was heuristically calculated by observing the time window between artifact and spike (bottom). **d)** Spikes from same neuron after artifact removal (top). Peri stimulus time histogram (PSTH) of neuron before and after correction of firing rate with artifact blanking method (bottom) Black bar indicates onset of stimulation. **e)** Peri-stimulus firing rates of M1, S1 and STR showed clear changes between early and late training sessions. Representative z-scored firing rates for one rat. (Red bar: M1 neurons, violet bar: S1 neurons and magenta bar: STR neurons, Brown bar: stimulation period). Time (0) corresponds to stimulation onset. **f)** Mean firing rate for the neuronal ensemble in ‘e’ showed clear differences between early (green) and late (blue) sessions. (Brown bar: stimulation period). **g,h)** To quantify changes in ensemble activity between early and late sessions, neuronal PSTHs were fed to a custom logistic regression classifier for predicting the stimulus type (‘virtual narrow’ vs ‘virtual wide’). Adaptive decoding i.e. retraining the classifier on each trial by updating the training set before making a prediction resulted in higher accuracy and lower mean error than non-adaptive decoding (n=6 rats). **i,j)** There was significant improvement in the decoding accuracy and mean error in late sessions as compared to early sessions (n=6 rats).

### Brain-To-Spine Interface

Once we established that rats can successfully discriminate DCS induced artificial somatic sensations and that these sensations can be decoded from neuronal firing patterns, we designed two experiments to test whether these artificial sensations can be transferred effectively between two rat brains, via a brain-to-spine interface (BTSI). In an initial experiment, we aimed to transfer natural tactile whisker responses from the brain of encoder rats to the spinal cord of decoder rats. Encoder rats were trained to discriminate between ‘narrow’ and ‘wide’ aperture widths using their large vibrissae in the aperture width tactile discrimination setup (Fig 1c and Fig 5a). Rats were rewarded for ‘narrow’ aperture widths in the left reward port and for ‘wide’ aperture widths in the right reward port.

**Figure 5:**
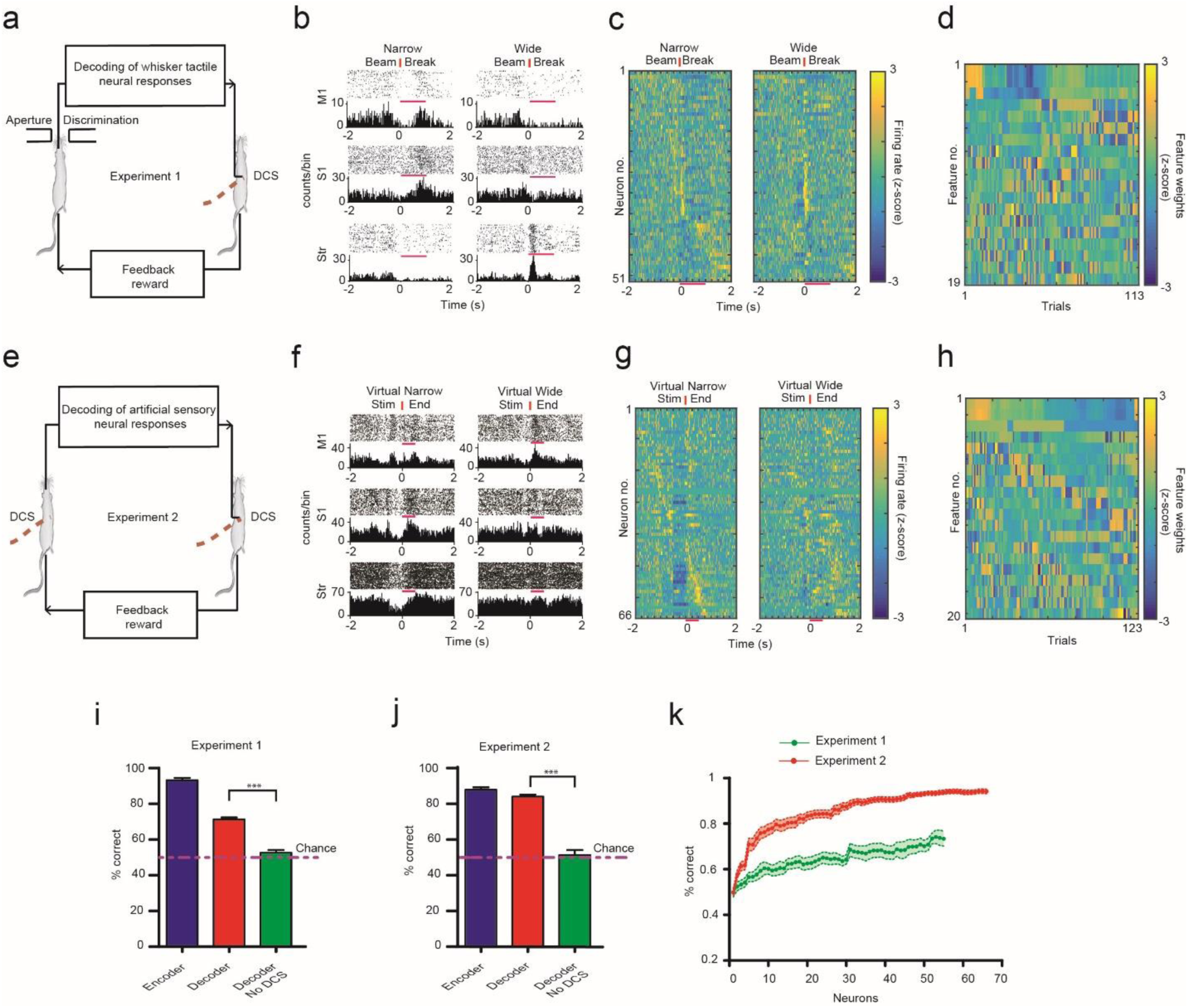
The Brain-to-Spine Interface. a) Schematic of the brain-to-spine-interface (BTSI) for Experiment 1 where neuronal activity from S1, M1, and STR areas of encoder’s brain during an aperture-width tactile discrimination task was transferred to spinal cord of the decoder. **b)** Raster plots and Peri-stimulus time histogram (PSTH) of representative neurons of encoder during ‘narrow’ and ‘wide’ aperture width trials. Time t=0 (red vertical line) corresponds to encoder rats crossing an infrared beam between the discrimination bars, indicating onset of ‘decoding epoch’ (pink lines above PSTHs) i.e. the neuronal activity from this time window was sent to the decoding algorithm **c)** Z-scored firing rates of all 51 neurons from one session rank ordered according to responses ± 2 seconds around beam break show differences between narrow and wide trials (Pink bars indicate decoding epoch). **d)** First 19 feature weights (z-scored) of the adaptive classifier can be seen changing throughout the session as the trials progressed. **e)** Schematic of the BTSI for Experiment 2 where neuronal activity from S1, M1, and St areas of encoder’s brain during DCS-induced sensory discrimination task was transferred to the spinal cord of decoder. **f)** Raster plots and PSTHs of representative neurons of encoder during ‘virtual narrow’ and ‘virtual wide’ trials with consistent labeling as in (b), except time t=0 indicates ‘end of stimulation’ and ‘decoding epoch’ (pink line) is 500 msec. **g)** Z-scored firing rates of all 66 neurons from one session with consistent labeling as in (c). **h)** First 20 feature weights (z-scored) can be seen changing throughout the session as the trials progressed. **i,j)** Performance of decoders during BTSI Experiment 1 (14 sessions, 6 decoders) and Experiment 2 (16 sessions, 6 decoders) was significantly higher than chance (p<0.001, Binomial test), while it dropped to chance levels when stimulation amplifiers were switched off in 6 sessions for each experiment type (***: p<0.001, Mann Whitney test). **k)** Neuron dropping curves for BTSI Experiment one (green, 14 sessions) and two (red, 16 sessions). Decoding accuracy of the classifier used for the logistic transformation dropped to chance levels (50%) as neurons were randomly removed from the ensemble.

As the encoder rats performed the whisker tactile discrimination task, single and multi-unit activity from M1, S1 and STR (total of 798 units from four encoder rats) was transformed into trains of electrical pulses using our custom adaptive logistic regression neural decoding classifier (see Methods for details). Then, these electrical pulses were sent to the spinal cord of decoder rats (n=6) which were trained to discriminate artificial sensations (schematic in Fig 5a left). Such functional link between encoder-decoder rat pairs defined the operation of a real-time BTSI. As expected, the electrical activity of M1, S1 and STR neurons, depicted by peri-stimulus time histogram (PSTH), exhibited considerable differences during narrow and wide trials (Fig 5b and 5c). Neuronal activity from the first 20 trials were used as input features to train an adaptive logistic regression classifier, which computed the probability of predicting a ‘narrow’ trial (a value between 0 and 1). Because of the adaptive nature of the decoding algorithm, from trial 21 onwards, PSTHs from all previous trials were added to the training set and the classifier was retrained for each subsequent trial, thus updating the feature weights used in the prediction. (Fig 5d). The number of pulses delivered to the decoder rat took values between 1 and 101, depending on the neuronal activity generated by the encoder’s brain. If decoder rats correctly replicated the behavior of the encoders, by perceiving the stimulation pattern as either ‘virtual narrow’ or ‘virtual wide’, they received a water reward. At the same time, encoders received an additional feedback reward. As expected, the encoder rats performed better (93.21 ± 1.33 % correct trials, Fig 4b) than the decoder rats (71.33 ± 1.04 % correct trials, Fig 5i) during the testing of this BTSI interface (n=14 sessions with 71.50 ± 4.48 trials per session). Although the performance of decoders was lower than encoders, it was considerably above chance levels (Binomial test: P<0.001 in all sessions), with the best session performance of 82% in 100 trials. As a control, we carried out the same BTSI experiment when the stimulation amplifiers used to deliver pulses to the decoder’s spinal cord were turned off. Under these conditions, the performance of the decoders was significantly reduced to chance levels (52.63 ± 1.48 % correct trials, p<0.001, Mann Whitney test, two-tailed, Fig 5i).

In our second experiment, the nature of the BTSI was similar to Experiment 1, except that decoding was performed on the neuronal response to artificial sensations created by DCS in the encoder rats, while they performed an artificial sensory discrimination task (Fig 5e). Both encoders and decoders were rats that had previously learned to successfully discriminate DCS patterns, except that the encoder rats in this experiment had received multi-electrode array implants in M1, S1, and Str, in addition to DCS electrodes. During the BTSI, the encoder rats received DCS with 51 pulses for the ‘virtual narrow’ trial and 1 pulse for ‘virtual wide’ trial. PSTHs describing ‘virtual narrow’ and ‘virtual wide’ trials were clearly different (Fig 5f and 5g). To eliminate the effect of the electrical stimulation artifact, only neuronal activity generated after the end of the electrical stimulation period was used to train the decoding classifiers. As the encoders performed an artificial sensory discrimination task, post-stimulus (500 msec after stimulation end) single and multi-unit activity (total of 1022 units from three encoder rats) was sent to our adaptive decoding classifier (classifier was retrained on each trial, see feature weights changing in Fig 5h) and transformed into DCS pulses, between 1 and 101, to be delivered to the spinal cord of decoder (n=6) rats. Encoder rats performed slightly better (87.88 ± 1.34 % correct trials, Fig 5j) than decoders (84.03 ± 1.21 % correct trials, Fig 5j) during operation of this BTSI interface (n=16 sessions with 97.75 ± 2.82 trials per session). The performance of decoders was considerably above chance levels (Binomial test: P<0.001 in all sessions), with best session performance of 92.22% in 90 trials. Similar to Experiment 1, the performance of decoders was significantly reduced to chance levels (51.37 ± 2.79 % correct trials, p<0.001, Mann Whitney test, two-tailed, Fig 5j) when the stimulation amplifier used for delivering the DCS to decoder animals was switched off.

We also investigated how our adaptive decoding classifier depended on the number of M1, S1 and STR neurons recorded simultaneously. Figure 5k illustrates the neuron dropping curves for both experiments 1 and 2, suggesting that neuronal ensembles with larger number of neurons outperformed those with smaller neurons in predicting the stimulus delivered.

The performance of decoder rats depended on how precisely the neuronal activity of encoders was decoded and transformed into a number of DCS pulses. Specifically, decoder rats performed an analog-to-digital (ADC) conversion; where analog input signal from the encoder (stimulation pulses 1 to 101) was converted into digital output choices (left: 1 vs right: 0) by the decoder. Fig 6a and 6b demonstrate decoder rats’ choice of left (virtual narrow) vs right (virtual wide) response for different DCS pulses delivered during a representative BTSI session of Experiments 1 and 2 respectively. Psychometric curves showing probability of left response as a function of DCS pulses illustrates clear trends in the decoder rats’ performance (Fig 6c). For example, the threshold for response was skewed to the left during Experiment 1 (21.49 pulses) as well as Experiment 2 (22.12 pulses). However, throughout the experimental session, decoder rats performed better in Experiment 2 as compared to Experiment 1 (Fig 6d). Overall, the decoder’s performance was clearly dependent on the neuronal representation of real or artificial tactile sensations generated by the encoder rat’s brain. This became evident by the discovery of a significant negative correlation between the decoder’s percent of correct trial discrimination and the mean squared error of the decoding classifier used for the logistic transformation of encoders’ neuronal activity across Experiments 1 and 2 (Fig 6e) (Spearman test: r= −0.8284, p<0.0001, two-tailed).

**Figure 6:**
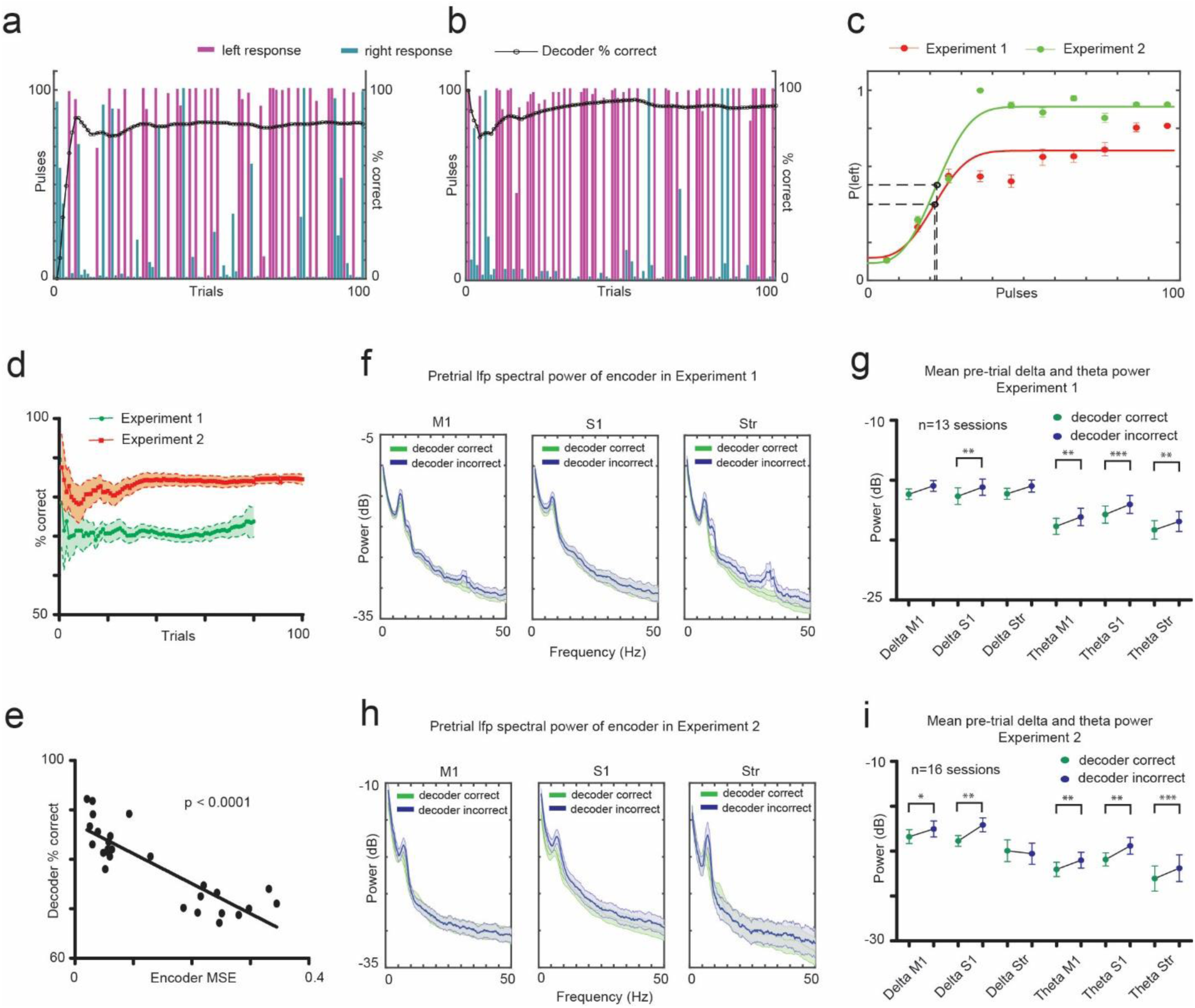
Dynamic nature of the Brain-to-Spine Interface. a) Pulses transformed from encoder’s neuronal activity that resulted in left response and right response by the decoder in a representative session from Experiment 1. **b)** Pulses that resulted in left response and right response of the decoder in a representative session from Experiment 2. Black circles in (a) and (b) indicate the performance of decoder in the session as a function of trial number. **c)** Probability of left response of decoder rats as a function of DCS pulses for all sessions of Experiment 1 (red, 14 sessions) and two (green, 16 sessions). Black circles indicate threshold number of pulses at inflection point of the curves. **d)** Mean performance of decoders in all sessions of Experiments 1 and 2 as a function of trial number indicates how the performance of the interface stabilizes as trials proceed. **e)** Performance of decoder showed significant negative correlation with the mean square error of the decoding classifier used to decode encoder’s ensemble neuronal activity (p<0.0001). **f,h)** Pre-trial LFP oscillatory power in M1, S1 and STR areas of encoder’s brain following correct or incorrect response from decoder in Experiments 1 and 2. Notice the clear difference in theta (5-9.5 Hz) frequency range **g,i)** Mean Pre-trial LFP oscillatory power for all sessions in the theta range calculated from M1, S1 and STR areas of encoder’s brain was significantly higher after decoder’s incorrect response than correct response for both experiments 1 and 2 (*: p<0.05, **: p<0.01, ***: p<0.001). Mean ± s.e.m.

Thereafter, we investigated the influence of the feedback reward provided by the decoder on the encoder’s brain activity as it prepared for the next trial. We compared the pre-trial (3000 msec) LFP activity in M1, S1 and STR areas of the encoders, after they had received a feedback reward on a previous trial, to the same brain area activity when it had not received such a feedback reward. We observed clear differences in spectral power in the low frequency bands (Fig 6f, 6h). Encoders’ mean pre-trial MI, S1 and STR LFP spectral power in the theta (5 – 9.5 Hz) band was significantly higher after they didn’t receive a feedback reward from the decoder, as compared to when they did receive an extra reward during both Experiments 1 and 2 (paired t-test, two-tailed, Experiment 1: 13 sessions, Experiment 2: 16 sessions, Fig 6g, 6i). This suggests that successful operation of a BTSI was influenced not only by how well the encoder rats encoded a given stimulus, and how well decoder rats were able to make sense of the information they received from the encoders, but also by how the encoder’s pre-trial cortical and striatal neuronal activity was modulated by the performance of their decoder counterparts during the previous trial. Put in another way, these findings, which reproduce and expand our previous observations using a BTBI that involved a cortical-to-cortical interface between encode-decoder rat dyads, clearly indicate that the optimal operation of a BTSI also involved bi-directional interactions between the encode-decoder rat dyads. Such a finding supports the hypothesis that, like BTBIs, BTSIs represent an interdependent behavioral-physiological computing system, comprising encoder-decoder dyads that take full advantage of using DCS as a channel for transmission of artificially-generated sensory signals to the brains of animals.

## Discussion

In the present study, we successfully trained rats to discriminate artificial tactile/somatic sensations created by temporally-varying DCS patterns. We demonstrated that learning to discriminate DCS-induced artificial sensations resulted in behavioral changes which led to cortical and striatal plasticity, as measured by both LFP and neuronal ensemble analysis. Finally, we developed a real-time brain-to-spine-interface where decoder rats successfully replicated the behavior of encoder rats relying on the transformation of the encoder rats’ neuronal activity into DCS patterns. Overall, we established that DCS can be used as a tool for transmission of prosthetic sensory information to the brain as well as for sensory transfer between multiple brains.

Monkeys and humans can discriminate artificial proprioceptive and tactile percepts generated by ICMS in the somatosensory cortex, while thalamic nuclei stimulation has been used to induce somatotopically organized percepts in humans^11, 14, 17, 18, 39, 40, 41, 42, 43^. However, to our knowledge this is the first time that DCS was used to generate artificial sensations to guide goal directed behavior in animals. In all, our rats were able to discriminate six unique pairs of stimuli that varied in time, but were delivered at constant amplitude (Fig 2a, Supplementary Fig 1a and 1b). Previous studies have shown discrimination between two patterns, which varied either spatially, or temporally, or spatiotemporally, given chance levels of 50%. In this study, we demonstrated that rats can consistently discriminate three (and likely four) distinct stimulation patterns within a session, which allowed us to reduce the chance levels to 33% and 25% respectively (Fig 2c, Supplementary Fig 1c). This is particularly important because DCS could be used as a carrier of sensory information to encode a vast array of tactile and proprioceptive sensations in patients with spinal cord injury or limb amputation.

In addition, we reported changes in striatal LFP spectral power in the delta and theta frequency ranges between early and late phases of DCS discrimination training. Incidentally, task related oscillatory peaks in delta and theta frequency ranges have been reported in striatal LFPs as rats performed a T-maze task^44^. The associative and sensorimotor regions of the striatum are modulated differently during early and late phases of skill learning and are known to be entrained differently to the theta band oscillatory rhythms^45, 46^. Moreover, cortical and striatal theta oscillations are known to modulate gamma power in cortical, striatal and hippocampal circuits^47, 48, 49^. These learning related modulations could explain the clear changes we observed, in terms of low frequency striatal spectral power, when we compared early and late trials during the learning of the discrimination tasks described here. Although we didn’t investigate M1-S1-STR functional connectivity, transcranial direct current stimulation and optogenetic stimulation of cortical structures has been reported to result in functional connectivity changes across multiple brain areas^50, 51^. By examining the M1, S1, and STR ensemble activity for stimulus prediction, we found significant improvements between early and late phases. Indeed, task related modulations in M1 and STR neurons have been observed in learning motor as well as abstract neuroprosthetic skills^38, 52, 53^. Future studies could investigate the role of individual cortical and striatal areas in facilitation of DCS-induced learning by using pharmacological inactivation or optogenetic inhibition of structures along the somatosensory pathway.

In this study, we observed a higher mean performance for decoder rats in comparison to the brain-to-brain interface (BTBI) study where sensorimotor information was transferred between the cortices of encoder-decoder rats using ICMS^32^. Moreover, the highest decoder rat performance (92%) obtained in this BTSI was greater than the maximum we achieved in both the ‘brain to brain’ (72%) as well as ‘brainet’ (87%) rodent studies^31, 32^. This suggests that DCS could be a viable alternative to ICMS as a method for delivery of sensory information. We observed that during the BTSI, the decoder’s performance in the sensory discrimination was negatively correlated with the mean squared error of the prediction of the original stimuli obtained from the encoder’s brain activity. Moreover, we also observed that low-frequency LPF oscillations in the encoders’ brain were significantly modulated by the feedback reward sent by the decoders. Incidentally, M1, S1 and dorsal premotor (pMd) neurons in primates increase their firing rates when reward expectation is not met^54, 55^. A recent study showed that M1 LFP spectral power in the 8-14 Hz band increased in non-rewarding trials as compared to rewarded trials in a primate center-out reaching task^56^. This is consistent with the increase in spectral power observed when encoder rats didn’t receive a feedback reward in our study. Thus the decoders’ performance in the BTSI was capable of influencing the encoders’ brain activity via a feedback reward. Therefore, due to the closed-loop nature of the BTSI, a clear sign of interdependence between encoder and decoder rats was established. This was precisely what we observed previously when rat pairs interacted through a BTBI^32^.

Decoder rats were initially trained to discriminate 1 pulse stimuli from 101 pulse stimuli, however once introduced to the BTSI, they had to convert continuous input data (any value between 1 and 101 pulses) into discrete output choices (virtual narrow vs virtual wide), essentially performing an analog-to-digital conversion. Decoder rats maintained the expected psychophysical profile, but the sigmoid curve was skewed towards the left (stimulation threshold = 21 or 22 pulses) due to the complex nature of the interface (Fig 6c). This was similar to our previous BTBI study where the stimulation response threshold was between 26-40 pulses^32^. Though the BTSI transfer bandwidth was 1 bit, we propose that it could be quickly scaled to 2 or 4 or 8 bits. This could be achieved by using spatio-temporal DCS patterns, spread across multiple electrode locations on the dorsal spinal cord, and by utilizing the organic computing framework highlighted in our previous work^31^.

In conclusion, we have demonstrated that DCS could be employed as a very optimal tool for delivering prosthetic sensory information to the brain within individual animals as well as between pairs of animals. In addition, we propose that DCS, a semi-invasive technique, could provide an alternative for cortical and subcortical stimulation to deliver tactile and proprioceptive feedback generated by a variety of neuroprosthetic devices, including those that are driven by brain-machine interfaces. Moreover, in the future, one could conceive the idea that BTSIs may be incorporated as part of new neuro-rehabilitation protocols aimed at restoring neurological functions in severely disabled patients. According to this view, non-invasive BTBI or BTSIs could be employed to directly link the brains of rehabilitation professionals and severely paralyzed patients in order to accelerate the process of cortical and subcortical plasticity needed for driving the patient’s functional recovery. In this approach, we envision that the therapist’s brain activity could be used as a source of feedback cues delivered to the patient’s spinal cord during the execution of a variety of sensory discrimination and motor tasks. Thus, through either a BTBI or a BTSI, a therapist-patient dyad could serve as the basis of a highly cooperative strategy that becomes incorporated into the patient’s long-term rehabilitation training protocol. Accordingly, we envision that DCS may play an important role in the functional rehabilitation of patients suffering from spinal cord injuries, Parkinson’s disease and, potentially, a variety of other neurological disorders (such as untreatable chronic epilepsy and other brain diseases that are mediated by localized subcortical seizures), and even possibly, psychiatric disorders, such as chronic untreatable depression.

## Methods

All animal procedures were performed according to prior approved protocols by the Duke University Institutional Animal Care and Use Committee and in accordance with National Institute of Health Guide for the Care and Use of Laboratory Animals. Long Evans rats weighing between 250-350 g were used in all experiments.

### Aperture width tactile discrimination task

Moderately water deprived rats were trained to perform a reinforcement learning based aperture width discrimination task after basic behavioral training as explained previously^35^. The task involved discrimination of narrow and wide aperture widths to receive a water reward. Rats were placed in the outer chamber and waited for the door to open to enter the inner chamber (see Fig 1c encoder section for schematic). Once inside, the rats had to pass through discrimination bars with adjustable widths in order to nose poke in the central wall. After a center nose poke, two reward doors opened in the outer chamber, which the rats had to lick in order to receive a water reward. Depending on the width of aperture, rats had to go to either port to receive 50ul of water reward. Narrow aperture width was rewarded in the left reward port while wide aperture in the right reward port. Incorrect responses were not rewarded. Tactile discrimination was assessed based on the percentage of correct trials. Some rats (n=4) that were trained on this tactile discrimination task, were then implanted with neural electrodes and used as encoders in Experiment 1 of the brain-to-spine interface (BTSI).

### Artificial sensory discrimination using DCS

A modified version of the aperture width tactile discrimination chamber explained above was used for training moderately water deprived rats (n=12) to discriminate DCS induced sensations except that the discrimination bars weren’t used and pulled back to maximum width (Fig 1b). After undergoing basic training that included: entering inner chamber, center poke in front wall, and poking left or right reward ports in the outer chamber, rats were trained to discriminate between a train of 100 pulses delivered at 100/333 Hz associated with a correct response in left reward port (Port 1 in Fig 1b) and 1 pulse corresponding to a correct response in right reward port (Port 2 in Fig 1b). DCS was initiated only when rats crossed the photobeam in the center of the inner chamber on their way to the center poke. Artificial sensations induced by 100 pulses and 1 pulse were named ‘Virtual narrow’ and ‘Virtual wide’ respectively. Correct responses in respective reward ports were rewarded with 50ul water. After ∼10 training sessions consistent performance of >80% was achieved (Fig 2a). In seven rats, single and multi-unit spikes as well as local field potential signals were recorded from multiple areas [(five rats: M1, S1 and STR); (one rat: M1 and S1); (1 rat: S1)] as they learned to discriminate DCS patterns over 11 sessions. Seven of the 12 rats trained with DCS were previously trained on the aperture width discrimination task. Three rats were naïve i.e. previously untrained on the aperture width task. Behavioral latency data in two rats wasn’t recorded due to technical limitations. Behavioral latency was quantified as time between inner chamber beam break and center poke (Fig 3a) Out of the twelve rats, those without brain implants were used as decoders in Experiment 1 of the BTSI, while those with brain implants were used as both encoders and decoders in Experiment 2 of the BTSI.

### Discriminating 3 and 4 artificial sensations induced by DCS patterns

Two rats that were trained to discriminate stimulation patterns associated with ‘virtual narrow’ (100 pulses at 333 Hz) and ‘virtual wide’ (1 pulse) sensations were further trained to discriminate three different DCS patterns. An additional reward port (port 3 in Fig 1b) was installed in the behavioral box that corresponded with the 3^rd^ stimulation pattern (100 pulses at 100 Hz). Initially, rats failed to associate the 3^rd^ reward port with 3^rd^ stimulus pattern, thus, to habituate them with the new reward port, we used a unique training paradigm (Supplementary Fig 1a). First, the novel stimuli was associated with port 3 by rewarding the rats when they made a response in the novel reward port. Once the rats successfully learned to associate the novel stimulation pattern with the new reward port, previously known stimulation patterns were paired with the 3rd stimuli (two unique combinations, stage II and III). After the rats reached performance criteria (>60%) in stages II and III, all three stimuli were simultaneously introduced in the same session to begin the three stimuli discrimination task (Fig 2c). Performance was analyzed based on the percentage of correct trials for the entire session and also the percent correct trials for each stimuli type.

Thereafter, one rat that still retained its spinal implant was trained to discriminate four different DCS patterns (4th stimulus pattern consisting of 100 pulses delivered in five bursts of 20 pulses each with inter-burst frequency of 2 Hz and inter pulse frequency of 333 Hz was associated with port 4 in Fig 1b). Similar to the three-stimuli training paradigm, initially, the 4^th^ novel stimuli was associated with port 4 in stage I, followed by three unique combinations of previously known stimuli with the 4^th^ stimuli, stages II, III and IV (Supplementary Fig 1b). Once the rat performed at criterion (>60%) at each stage, the four-stimuli discrimination task began (Supplementary Fig 1c). Performance was analyzed based on percentage of correct trials for entire session and also percent correct trials for each stimuli type.

Irrespective of the stage of learning (introduction to novel stimuli or discriminating pairs of stimulus combinations or all stimuli together), chance performance was maintained at either 33% (learning three stimuli) or 25% (learning four stimuli) by opening all reward ports simultaneously after the center nose poke, thereby minimizing the probability of chance response. This unique paradigm made the learning task harder for the rats. Discrimination performance plots for each stimuli type were generated by selecting trials belonging to that particular stimuli within the session and calculating the percentage of correct responses on those trials (Figs 2b, 2d, and Supplementary Fig 1d). To better understand the response choices for each stimuli type, we first classified the trials based on stimuli type, then among those trials identified the responses chosen, and plotted them as percentages for each session during the training period (Supplementary Figs 1e and 1f). This confirmed that as the training period progressed response choice percentages associated with the particular stimuli type (e.g. response 1 for stimuli 1) increased gradually.

### Brain-to-Spine Interface

In Experiment 1, as encoder rats performed whisker tactile discrimination task, their cortical and sub-cortical (M1, S1 and STR) single and multi-unit activity was sent to a neural decoder (explained below) which transformed it into the number of DCS pulses to be applied to the spinal cord of decoder rats (Figs 1c and 5a). The first 20 trials of the interface were used to create a training set for the neural decoding algorithm. During this phase the number of pulses delivered to the decoder were 1 for a ‘wide width’ trial and 101 for a ‘narrow width’ trial. From trial 21 onwards, the number of DCS pulses sent to the decoder rat (varying between 1 and 101 pulses) were calculated by transforming the corticostriatal ensemble activity of the encoder rat. If the decoder rat replicated encoder’s response choice correctly, then both rats received a water reward (this constituted as additional feedback reward for the encoder). The decoder’s performance (Fig 5i) depended not only on the classification accuracy of encoder’s ensemble activity (whether the neural activity predicted encoder’s response correctly) but also on its prediction error (how far the value of pulses delivered was from either 1 or 101).

In Experiment 2, rats that had already learned to discriminate two DCS patterns were used as encoders and decoders (Figs 1c and 5e). Rats that had neuronal as well as spinal implants were used as encoders while those with only spinal implants were used as decoders. Encoder rats received 101 pulses for ‘virtual narrow’ and 1 pulse for ‘virtual wide’ trials. Post-stimulus single and multi-unit ensemble activity from M1, S1 and STR of encoder rats during artificial sensory discrimination was transformed by the neural decoding algorithm into pulses that ranged from 1 to 101. These pulses were delivered to the decoder rats using DCS. Similar to Experiment 1, if the decoder performed correctly, both rats received a water reward (constituting additional feedback reward for encoder).

### Neural decoding algorithm

Single and multi-unit activity was sampled for 1 sec starting from the time the encoder rat crossed the photobeam inside the inner chamber for Experiment 1 and for 500 msec starting from the time stimulation ended (to avoid effect of stimulation artifact) for encoder rat in Experiment 2. Spiking data was binned in 250 msec and 100 msec bins respectively for Experiments 1 and 2. The initial training set was created using the first 20 trials of the encoder, where each data bin of each neuron was considered as a feature, and used to predict the number of pulses by training a custom adaptive logistic regression machine learning algorithm. Cost function to be minimized for the learning algorithm was computed using the equation,

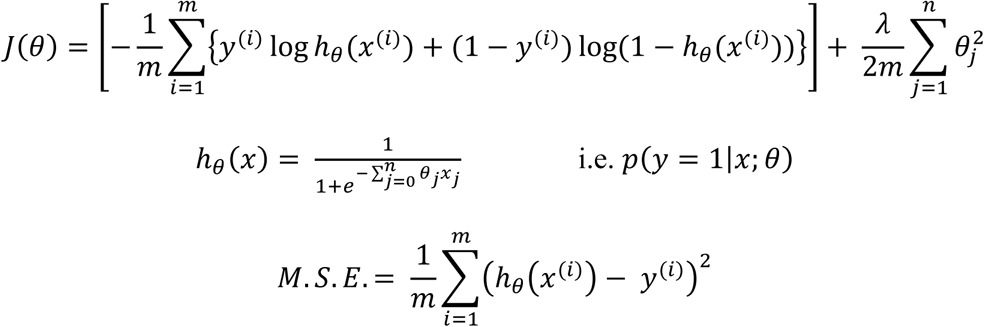

Where, *x*^(*i*)^ contains all binned firing rates of all neurons called as ‘features’ (‘n’ in number) for trial (*i*), *y*^(*i*)^ is 1 for narrow trial and 0 for wide trial, *θ_j_* is the feature weight corresponding to the *j^th^* feature. *h_θ_*(*x*) was the logistic transformation that computed the probability of predicting a narrow trial. If *h_θ_*(*x*) was greater than or equal to 0.5, the algorithm was said to predict a narrow trial. Dimensionality of features was reduced such that 99% of variance was retained in the training data set by using the ‘SVD’ function. ‘A ‘gradient descent’ optimization approach with regularization parameter λ set at 1 was implemented using the ‘fminunc’ function in order to minimize the regression cost function *J*(*θ*).

From trial 21 onwards, as the brain-to-spine interface began, binned firing rates of encoder neurons from all previous trials were included to create an updated training set which was used to re-train the classifier and make a prediction for the current trial. Because the classifier was updated on every single trial based on data from all previous trials, the logistic transformation feature weights θ changed throughout the session (Figs 5d and 5h).

The above logistic regression classifier was also used for classifying stimuli type based on neuronal ensemble activity in rats that learned artificial sensory discrimination. There, we evaluated the performance of the decoding algorithm by measuring its decoding accuracy and mean squared error. We observed that its performance was better in the adaptive version i.e. when the classifier was retrained on each trial as compared to the non-adaptive version i.e. when the classifier was only trained once using first 20 trials (Figs 4g and 4h). All scripts were custom written in Matlab (8.4.0, Mathworks, Natick, MA) using in-house functions. Decoding accuracy was defined as the percentage of trials that the algorithm predicted correctly (Figs 4g, 4i and 6e). Prediction error of the algorithm was defined as the mean squared error (M.S.E.) of *h_θ_*(*x*) calculated for all trials of the session (Figs 4h, 4j and 6e). To quantify neuronal plasticity between early and late training sessions, we compared decoding accuracy and MSE between those sessions.

Neuron dropping curves were generated from encoder’s ensemble activity by calculating prediction accuracy of the decoding algorithm for each BTSI session by randomly dropping neurons from the ensemble until only one neuron was left (Fig 5k).

### Surgery for implantation of DCS electrode and microelectrode array

Custom built movable microelectrode bundles or fixed arrays were implanted in the M1, S1 and striatal areas (STR) of rats (Fig 1a). The stereotaxic coordinates for each of the areas relative to bregma are: M1 [AP: 0 mm; ML: 1.6 mm; DV: −1 mm], S1 [AP: −3 mm; ML: 5.5 mm; DV: −1.25 mm], STR [AP: 0 mm; ML: 3.5 mm; DV: −4.5 mm]. Rats that were trained on the aperture width discrimination task and used as an encoder for the BTSI Experiment 1 (n=4) were implanted in M1, S1 and STR. Out of all rats that were trained to discriminate artificial sensations generated using DCS, seven rats had recording microelectrodes. From these, five rats had electrodes in M1, S1 and STR, one rat had electrodes in M1 and S1 while one rat had electrodes only in S1. Three of these seven rats were used as encoders for BTSI Experiment 2. Bipolar dorsal column stimulation electrodes were implanted in the epidural space between T2 vertebra and spinal cord as explained previously^29^.

### Neurophysiological recording

Neuronal single and multi-unit activity and Local Field Potential (LFP) data was recorded using a Multichannel Acquisition Processor (64 channels, Plexon Inc., Dallas, TX) as described previously^57^. Neural signals were differentially amplified (20,000 − 32,000X), filtered (400Hz – 5 kHz) and digitized at 40 kHz. LFPs were pre-amplified (1000X) and digitized at 1000 Hz. Single and multi-units were sorted online using (SortClient 2002, Plexon Inc., Dallas, TX) and interfaced with Matlab (8.4.0, Mathworks, Natick, MA) for real-time analysis and decoding.

In rats that learned to discriminate DCS patterns and those that had neural electrodes (n=7), to study the effect of DCS induced learning on the neuronal ensemble activity, we sorted the neuronal data from first and last learning sessions into single units using Offline Sorter (Version 2.8.8 Plexon Inc, Dallas, TX). One rat’s event data from the first session was missing and hence it was excluded from this analysis. We observed that stimulus artifact could be easily separated from the neuron cluster using principal components (Figs 4a and 4b). To compensate for effect of amplifier dead time during the artifact, we heuristically assigned a blanking period of ‘4ms’ after each stimulation pulse (Fig 4c), and recalculated the firing rates for each neuron (Fig 4d). These corrected firing rates were then used to generate PSTHs (Figs 4e and 4f), which were employed to train the neural decoding algorithm and calculate the effect of DCS learning on the decoding accuracy and mean squared error as explained above.

### Dorsal column stimulation

DCS consisted of charge balanced biphasic square pulses of 200 or 1000 us duration delivered at 100-333 Hz, either as ‘trains’ or ‘bursts’ of pulses depending on the condition of the experiment. Stimulation intensities were determined during each session and set at a value between sensory and discomfort threshold (therapeutic range with intensities that varied from 110 µA to 600 µA). Specific patterns of stimulation pulses were generated by custom scripts written in Matlab (8.4.0, Mathworks, Natick, MA) that controlled the electrical microstimulator (Master 8, AMPI, Jerusalem, Israel).

### LFP analysis

Raw LFP time series was standardized and stimulation artifacts were detected and indexed. Because LFPs during the stimulation periods were often saturated by the artifact, pre-stimulation and post-stimulation periods were used in the spectral analysis. Spectral power was computed using the multi-taper method with five tapers using Chronux 2.0^58^. For spectral analysis of the pre-stim period, 300 msec time segments before stimulation onset on each trial were collected to calculate the average spectral power using the ‘mtspectrumc’ function. Spectral power was log transformed into dB and averaged per area.

During analysis of pre-trial spectral power of encoder after decoder’s correct/incorrect response in the BTSI experiments, we observed that the correct response trials outnumbered incorrect response trials by a factor of 2-5. When we calculated pre-trial spectral power after correct decoder response by using randomly-selected subset of trials that matched the number of trials after incorrect decoder response, the results showed the same statistical significance compared to when we didn’t down-sample the trials. Thus, results in this paper contain the analysis using the actual number of trials even though the number of trials after decoder’s correct and incorrect response were different. Pre-trial encoder spectral power was calculated using ‘3000 msec’ time segments before trial onset using the ‘mtspectrumc’ function, log transformed into dB and averaged per area. Spectral power measures throughout this study were averaged in the following frequency bands for statistical analysis: delta (1.5 – 4.5 Hz), theta (5 – 9.5 Hz), beta (10 – 25 Hz), low gamma (26 − 40 Hz) and high gamma (65 – 99 Hz).

### Statistical Analysis

Percentage correct trials were used to assess behavioral performance of all rats during individual training as well as during the brain-to-spine interface. Performance of the decoding algorithm was measured by calculating the percentage of trials correctly predicted from the entire session. Prediction error was calculated as the mean squared error of actual value from the predicted value using all trials from the session.

Comparison between learning rates of individual stimulation patterns was performed using linear regression and checking for slopes. Comparison between ‘early’ and ‘late’ spectral power values, % correct trials, latency, and between pre-trial spectral powers after correct and incorrect decoder response during the BTSI, was performed using paired t-test or Wilcoxon signed rank test whenever the distributions failed the normality test. The Mann Whitney test was used to compare the performance of decoder during BTSI when the stimulation amplifier was ‘ON’ or ‘OFF’. Correlation between spectral power change and behavioral latency change was tested for significance using the Pearson test while correlation between decoder’s performance and mean squared error of encoder’s neural classifier was tested using the Spearman’s rank correlation test.

## Acknowledgements

We thank J. Meloy for help with building recording electrodes; L. Oliveira and S. Halkiotis for technical support, E. Thomson for comments, and M. Pais-Vieira for initial inspiration. This research was supported by NIH Transformative award (R01-NS073125-03) and NIH Director’s Pioneer Award (DP1-OD006798). The content is solely the responsibility of the authors and does not necessarily represent the official views of the Office of the NIH Director or the NIH.

## Author Contributions

A.P.Y. and M.A.L.N. designed experiments; A.P.Y. and M.A.L.N. wrote the paper; A.P.Y. and M.A.L.N analyzed the data; A.P.Y., and D.L. conducted experiments.

## Supplementary Information

**Supplementary Figure 1.**
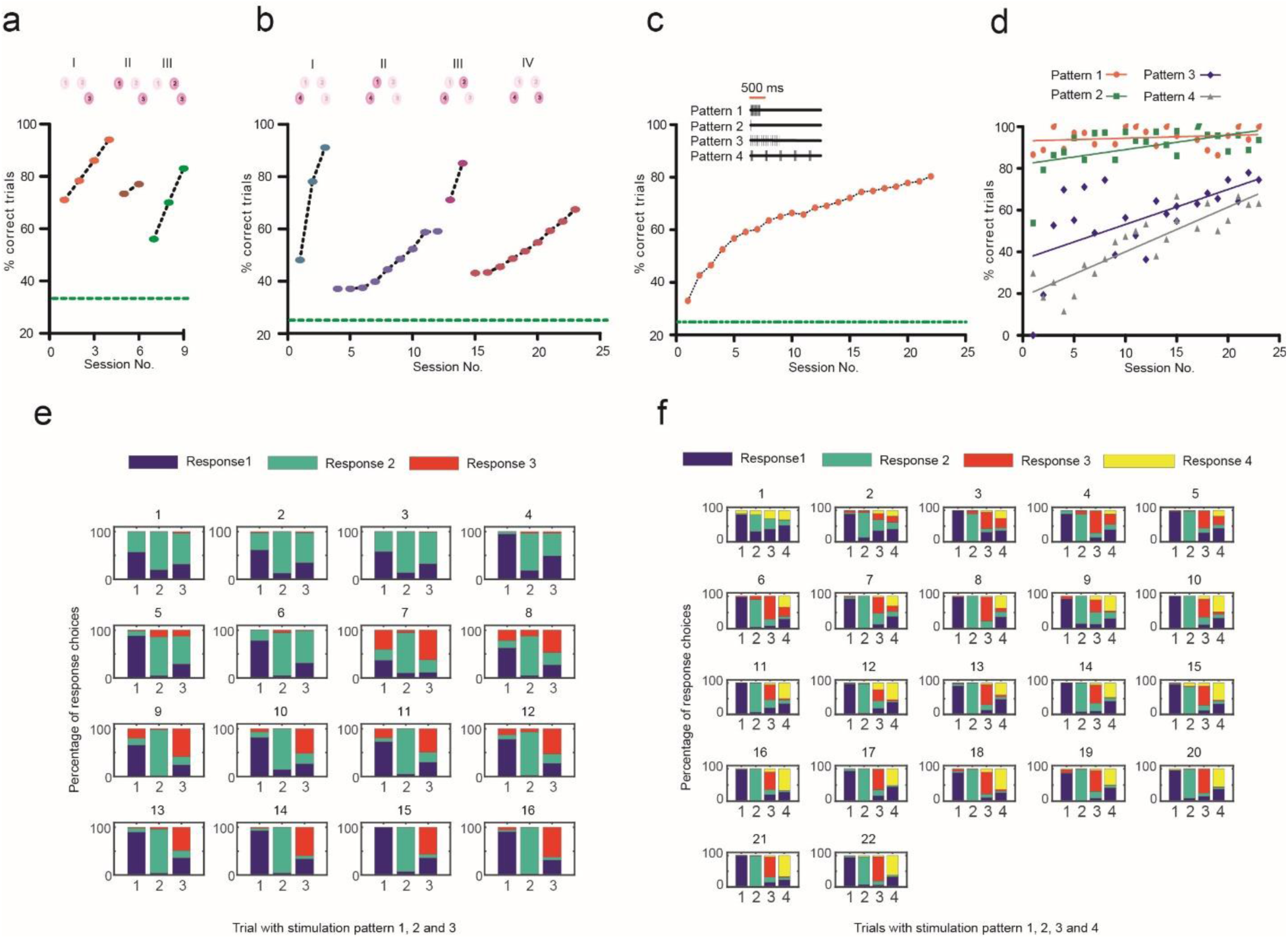
a) Training paradigm to associate novel stimulation pattern (Pattern 3) with reward port 3 for one rat. In stage I, all trials in the session belonged to Pattern 3 (100 pulses at 100Hz). During stages II and III, trials for both Pattern 3 and Pattern 1, and Pattern 3 and Pattern 2 were respectively applied in the training sessions. Once the rat performed >75% correct in stages II and III, training sessions with all three patterns together were initiated as shown in Fig 2c. Chance performance was kept at 33% during all stages by giving the rat access to make a nose poke at any of the three reward ports (green line). **b)** Training paradigm to associate novel stimulation pattern (Pattern 4) with reward port 4 for one rat. In stage I, all trials in the session belonged to Pattern 4 (100 pulses in 5 bursts of 20 pulses each with inter-burst frequency of 2 Hz and inter pulse frequency of 333 Hz). During stages II, III, and IV, trials for both Pattern 4 and Pattern 1, Pattern 4 and Pattern 2, and Pattern 4 and Pattern 3, were respectively applied in the training sessions. Once the rat performed >60% correct in stages II, III, and IV, training sessions with all four patterns together were initiated. Chance performance was kept at 25% during all stages by giving the rat access to make a nose poke at any of the four reward ports (green line). **c)** Learning curve showing one rat learned to discriminate 4 DCS patterns (Pattern 1: 100 pulses at 333 Hz, Pattern 2: 1 pulse, Pattern 3: 100 pulses at 100 Hz, and Pattern 4: 100 pulses in five bursts of 20 pulses each). Graph shows percentage correct trials for 22 sessions, green line represents chance level of 25 %). **d)** Discrimination performance for each of the four patterns shown individually across sessions. **e)** Percentage of response choices classified by trial type for one rat as it learned to discriminate three stimulation patterns. Stacked bar graphs are color coded for response type. Trial type is indicated on x-axis. Notice how % response 1 on trial type 1, % response 2 on trial type 2, and % response 3 on trial type 3, increases from session 1 to 16. **f)** Percentage of response choices classified by trial type for one rat as it learned to discriminate four stimulation patterns. Stacked bar graphs are color coded for response type. Trial type is indicated on x-axis. Notice how % response 1 on trial type 1, % response 2 on trial type 2, % response 3 on trial type 3, and % response on trial type 4, increases from session 1 to 22.

